# Variations in perfusion detectable in advance of microstructure in white matter aging

**DOI:** 10.1101/2023.06.30.547294

**Authors:** Tyler D. Robinson, Yutong L. Sun, Paul T. H. Chang, Claudine J. Gauthier, J. Jean Chen

## Abstract

One of the most promising interventional targets for brain health is cerebral perfusion, but its link to white matter (WM) aging remains unclear. Motivated by existing literature demonstrating links between declining cortical perfusion and the development of WM hyperintensities, we posit that regional WM hypoperfusion precedes deteriorating WM integrity. Using the Human Connectome Project Aging (HCP-A) data set, we examine tract-wise associations between WM microstructural integrity (i.e. fractional anisotropy, mean diffusivity, axial diffusivity, and radial diffusivity) and perfusion (i.e. cerebral blood flow and arterial transit time) in ten major bilateral WM tracts. Results show that tracts displaying the largest CBF decline in aging do not necessarily display the largest ATT decline, and vice versa. Moreover, significant WM perfusion-microstructure canonical correlations were found in all tracts, but the drivers of these correlations vary by both tract and sex, with female subjects demonstrating more tracts with large microstructural variations contributing to the correlations. Additionally, arterial transit time appears to be the earliest indicator of WM declines, preceding age-related microstructural differences and CBF in several tracts. This study contributes compelling evidence to the vascular hypothesis of WM degeneration, and highlights the utility of blood-flow timing as an early marker of aging.

## 1 INTRODUCTION

Increasing appreciation for the importance of cerebrovascular health in maintaining tissue integrity has led to the examination of tissue structure-perfusion associations in brain aging. The existing literature has focused largely on gray matter (GM) perfusion (Zhang et al., 2017) and its associations with cortical and subcortical atrophy (Baker et al., 2014; Chen et al., 2011, 2013; C.-M. Kim et al., 2020; Mayer et al., 2021). In the white matter (WM), microstructural declines in aging are reported in a robust body of literature, demonstrating regional differences in age-related microstructural declines along both broad regional and tract-specific segmentations (Baker et al., 2014; Bender et al., 2016; de Groot et al., 2015, 2016; Sala et al., 2012; Slater et al., 2019). Furthermore, recent investigation of tract-wise differences in patterns of microstructural and morphometric decline (Schilling et al., 2022), as well as in relationships between WM integrity and tract morphometry (Robinson et al., 2024), suggest that microstructural variations in different tracts are differentially associated with morphometric declines with age. However, the basis of this heterogeneity between tracts remains unclear. These findings cumulatively suggest that a more complete understanding of the interrelations between perfusion and integrity in the WM on a tract-specific basis is necessary to clarify patterns of decline across the brain.

A well-established relationship between cortical perfusion and the development of WM hyperintensities in associated subcortical regions illustrates the importance of CBF to maintaining the integrity of WM fibers (Bernbaum et al., 2015; O’Sullivan et al., 2002). Cortical perfusion was further related to subcortical WM integrity in normal aging (Chen et al., 2013). Relationships have been demonstrated between cortical perfusion and cognitive performance (D. Kim et al., 2021), and decreased cortical perfusion may predict heightened vulnerability to pathological declines (Chen et al., 2013; Mayer et al., 2021; O’Sullivan et al., 2002). However, regional variation in age-related cortical atrophy does not appear to match regional reductions in cerebral blood flow (CBF) (Chen et al., 2011; Tosun et al., 2010), suggesting a potential temporal asynchrony in perfusion and structural effects of age. Such an asynchrony, with perfusion decline preceding structural decline, would be consistent with the vascular hypothesis of aging and dementia (de la Torre, 2018), but has yet to be confirmed despite promising evidence that CBF changes precede structural degeneration changes in the GM (Lee, 2022; Wierenga et al., 2014; Zlokovic, 2011). Moreover, investigations of this nature are much scarcer in the WM, despite the latter manifesting age effects earlier than GM (Giorgio et al., 2010). Existing research has demonstrated that trends of WM perfusion variations with age may differ significantly from those found in the GM, to the extent that the direction of perfusion change in aging is debated (Alisch et al., 2021; Damestani et al., 2024). Clarifying these regional patterns by tract will be necessary to understanding perfusion as a potential precursor to microstructural declines.

If it can be established that WM perfusion changes similarly lead microstructural degeneration, then regional perfusion in the WM itself can be a strong candidate as an early imaging marker of age-related declines in WM microstructure should, but the structural-perfusion relationship in the aging WM has not been examined. This paucity of work on WM perfusion is largely due to challenges in WM perfusion quantification (Bernbaum et al., 2015; Chen et al., 2013), the low vascularization and long arterial transit delays (ATT) in the WM as well as the presence of partial-volume effects with the GM. Recently, multi-delay pseudo-continuous ASL, which is better suited for WM perfusion mapping than conventional techniques, was used to demonstrate tract-specific declines in CBF and elongation in ATT (Damestani et al., 2024). Building on this development and incorporating regional perfusion measures into our ongoing model of WM microstructural and morphometric declines with age (Robinson et al., 2024), we aim to gain a more comprehensive understanding of how WM microstructural variations across age are related to perfusion parameters. To some extent, patterns of WM decline and resilience appear to broadly follow developmental trajectories, though support for ordered models is mixed (Hoagey et al., 2019; Slater et al., 2019; Stricker et al., 2009), prompting a tract-wise approach. Given the previously limited ability to measure regional differences in WM perfusion (Bernbaum et al., 2015; Chen et al., 2013) and the sparsity of studies addressing WM vascular-microstructural declines on a tractwise basis (Mayer et al., 2021; Smirnov et al., 2021), this study aims to fill both of these knowledge gaps, with the understanding that WM tracts represent the meaningful structural units for segmentation of the WM (Baker et al., 2014; Sala et al., 2012; Yeatman et al., 2014) in aging, as well as reflecting functional connections between terminal cortical regions (Nozais et al., 2021). As variability in tractwise perfusion remains underexplored in healthy adults, it represents a natural extension to the existing literature and a valuable missing component in modeling regional WM declines with age and may represent an underlying source of differences in order and rate of tractwise declines. Using the Human Connectome Project Aging (HCP-A) data set (Bookheimer et al., 2019), this study sought to map tractwise age-related variations in WM perfusion, as well as the relationships between tractwise WM perfusion and microstructure. We hypothesized that (1) tract perfusion and microstructure will not decline at the same rate at aging, and that (2) tractwise perfusion changes will be detectable at younger ages than microstructural declines.

## 2 METHODS

This study includes five hundred and thirty-five healthy adult subjects (300 female, aged 36-100) from the Human Connectome Project in Aging (HCP-A) dataset (OMB Control# 0925-0667 (Bookheimer et al., 2019; Harms et al., 2018). All subjects were in generally good health and without pathological cognitive impairment (i.e. stroke, clinical dementia). Following manual quality control, thirty-two subjects were excluded from this sample due to MRI artifacts that produced notable errors when generating WM masks. Three additional subjects were excluded as age outliers due to being the only subjects of 100 years of age. Thus, the final sample size was 500 subjects (294 female, aged 36-89, aprox. 59% postmenopausal (Gold, 2011)). Hypertension status was determined based on mean arterial pressure (Kandil et al., 2023).

**Table 1:**
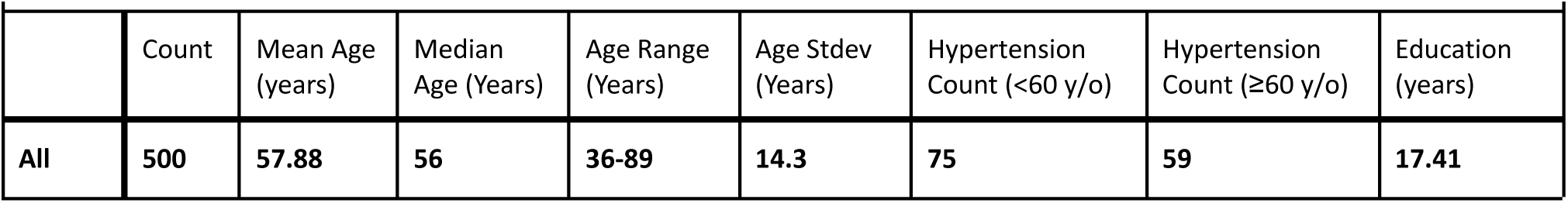
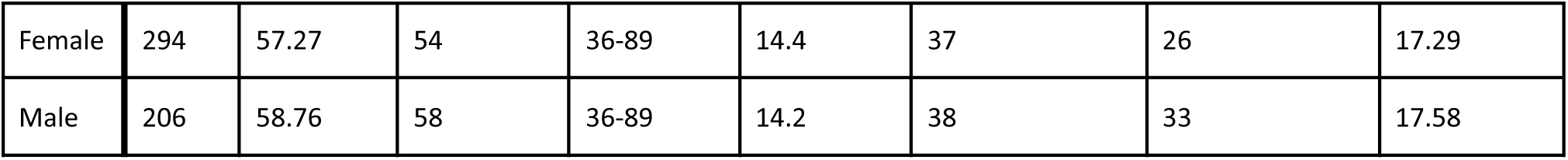
Sample Demographics Demographic statistics for all subjects, female subjects, and male subjects. All subjects were in good health and without pathological cognitive impairment (i.e. stroke, clinical dementia).

|  | Count | Mean Age (years) | Median Age (Years) | Age Range (Years) | Age Stdev (Years) | Hypertension Count (<60 y/o) | Hypertension Count (≥60 y/o) | Education (years) |
| --- | --- | --- | --- | --- | --- | --- | --- | --- |
| All | 500 | 57.88 | 56 | 36-89 | 14.3 | 75 | 59 | 17.41 |
| Female | 294 | 57.27 | 54 | 36-89 | 14.4 | 37 | 26 | 17.29 |
| Male | 206 | 58.76 | 58 | 36-89 | 14.2 | 38 | 33 | 17.58 |

The study accessed whole-brain T1-weighted structural MRI, diffusion-weighted MRI (dMRI) with a (1.5mm)^3^ voxel resolution, MB=4, with 93 directions at b=1500s/mm^2^, and multi-delay multi-band echo planar imaging (MB-EPI) pseudo-continuous arterial spin labeling (pCASL) with a (2.5mm)^3^ voxel resolution, MB=6, with TIs = 0.7, 1.2, 1.7, 2.2, and 2.7, collected using four matched Siemens Prisma 3T MRI scanners. Further imaging parameters for structural, diffusion, and perfusion imaging can be found in (Harms et al., 2018).

### 2.1 Image analysis

dMRI data were corrected for eddy-current and susceptibility-related distortions via EDDY prior to use in tractography analysis. Eighteen major WM tracts were reconstructed from the dMRI data using the Tracts Constrained by Underlying Anatomy (TRACULA) tool in Freesurfer version 7.2.0 (Maffei et al., 2021; Yendiki et al., 2011). Tracts reconstructed by TRACULA’s trac-all command represent pathways in which each adjacent voxel has a 99% probability of containing the next step in the fibre pathway of interest, ensuring the highest level of anatomical feasibility while excluding potential WM hyperintensities. Prior to extracting diffusion values, all values below 20% of the maximum posterior probability distribution were thresholded (Maffei et al., 2023). The TRACULA pipeline also incorporates correction for eddy current distortion, brain extraction, tensor calculation, and affine registration. Per-subject DTI and Perfusion images were registered in local space using Freesurfer’s TKRegister function. FSL’s fslmaths function was then used to produce per-subject masks of each WM tract in local space from the full TRACULA tract volumes. The eighteen reconstructed tracts were combined bilaterally to produce ten tracts of interest for analysis: The major forceps (Fmajor), minor forceps (Fminor), anterior thalamic radiation (ATR), cingulum angular bundle (CAB), cingulate gyrus (CCG), corticospinal tract (CST), inferior longitudinal fasciculus (ILF), superior longitudinal fasciculus parietal (SLFP), superior longitudinal fasciculus temporal (SLFT), and the uncinate fasciculus (UNC) (**Figure 1**). Rosner’s tests were performed to identify and exclude volume outliers per tract in single tract analyses. The resulting subject counts for these analyses were: Fmajor (N=493), Fminor (N=493), ATR (N=497), CAB (N=490), CCG (N=495), CST (N=491), ILF (N=497), SLFP (N=497), SLFT (N=495), and UNC (N=495).

**Figure 1:**
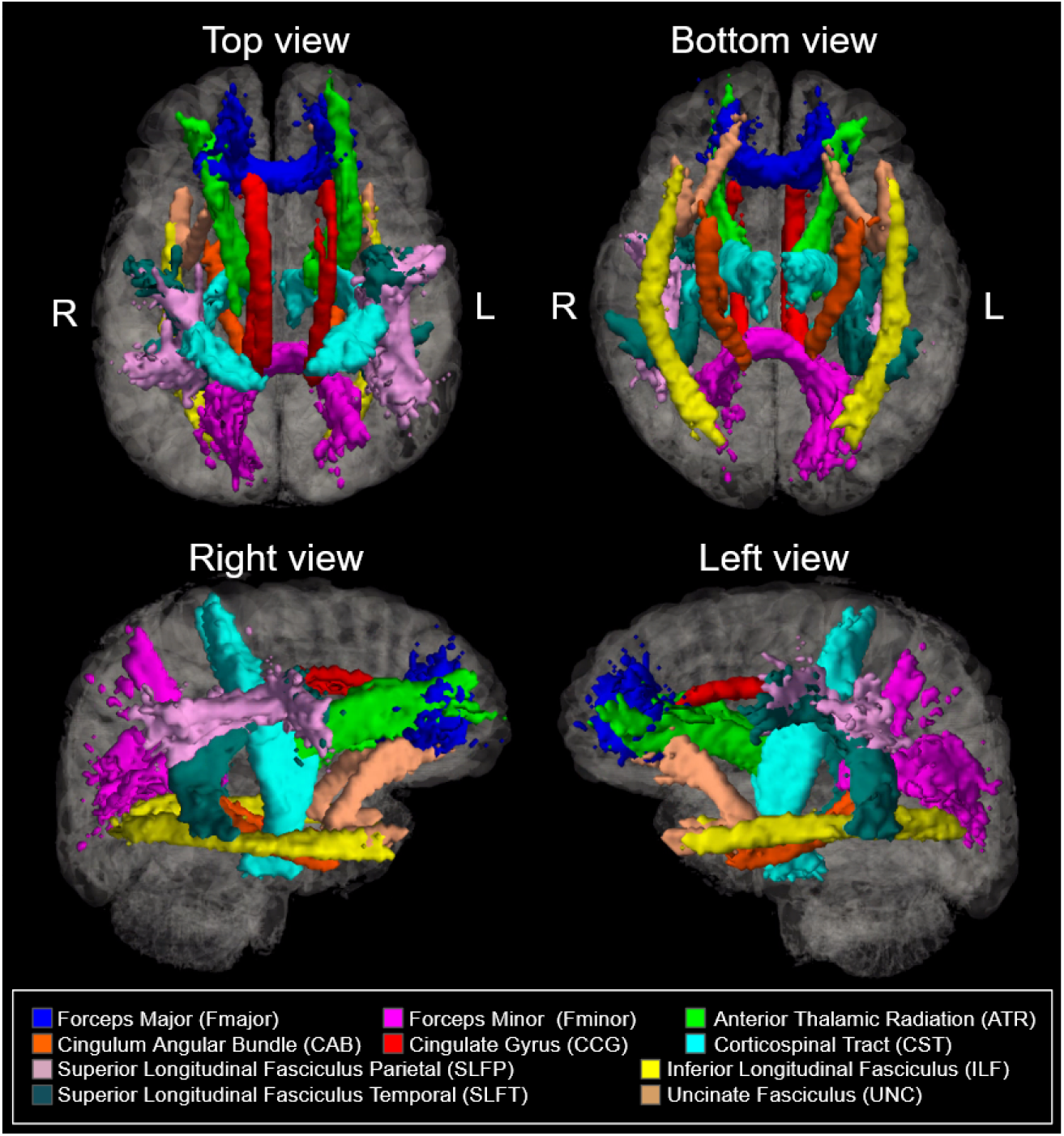
Ten bilateral tracts of interest reconstructed Freesurfer’s TRACULA package. Note that there is limited overlap between tracts.

#### 2.1.1 Microstructure

WM microstructural integrity was quantified using fractional anisotropy (FA), mean diffusivity (MD), axial diffusivity (AD), and radial diffusivity (RD). FA, MD, AD, and RD maps were derived via Dipy’s DKI script to achieve kurtosis-corrected DTI metric fitting, due to the high b-value used in the HCP-A acquisitions. TRACULA generated tract masks were applied to local dMRI images in flsmaths to generate tract means for FA, MD, AD, and RD. For each metric, we defined the “baseline” as being measured from the youngest 10% of the group, and computed baseline normalized (percent of baseline) values (“FA_perc_”, “MD_perc_”, “AD_perc_”, “RD_perc_”). These were defined as individual raw tract values divided by the mean raw tract value of these youngest subjects to assess percent differences from the youngest baseline, with separate baseline calculations performed for female and and male subsets of the sample (**Figure 2a**).

#### 2.1.2 Perfusion

Perfusion metrics of interest to this study included cerebral blood flow (CBF) and arterial transit time (ATT). A separate calibration scan was used to measure M0, which was used in CBF quantification. The pCASL label elevated the WM SNR relative to conventional pulsed ASL methods. Multi-TI data were fit to the general kinetic model (implemented through FSL’s oxford_ASL command) to produce local CBF and ATT maps (Wang et al., 2013). Partial volume correction was applied to the local maps. Specifically, the T1-weighted structural image for each subject was segmented, from which GM and WM partial volume estimates were extracted and transformed into the same space as the ASL images. GM and WM masks were generated using empirically chosen thresholds of 0.75 and 0.9, respectively, and applied to the estimated CBF images within which the mean estimated CBF and its standard deviation estimate were calculated. Separate gray and white matter T1s are used in the kinetic model inversion. Multi-tissue T1 values were adopted from the literature to account for partial-volume effects. Thus, the use of multi-TI pCASL with partial-volume correction addresses the challenges of long ATT, low SNR and partial-voluming in WM perfusion mapping.

As with microstructure, TRACULA-generated tract masks were applied to the local maps in fslmaths to produce tract means for both perfusion metrics. Baseline normalized perfusion values (“CBF_perc_”, “ATT_perc_”) were again calculated as individual raw tract values divided by the mean raw tract value of the youngest 10% of subjects (i.e. the baseline group) using the same female and male subsets as previous baseline.

### 2.2 Statistical analysis

#### 2.2.1 Age and sex effects

Between-subjects ANOVA was used to determine the presence of sex differences in tract specific FA_perc_, MD_perc_, CBF_perc_, and ATT_perc_. Subsequent assessment of sex differences in microstructure and perfusion were conducted for each tract via sequential multivariate regressions of sex on to mean FA_perc_, MD_perc_, CBF_perc_, and ATT_perc_ using age as a covariate of no interest. Whole-WM age-related microstructure and perfusion differences were assessed via multivariate quadratic regression by first regressing age onto mean FA_perc_, MD_perc_,CBF_perc_, and ATT_perc_ values derived from voxel-count weighted tract averages using sex as a covariate of no interest. This process was repeated for each tract separately in tract-wise analyses. FDR adjustment was again applied to the resulting p-values. Sex differences in age effects on tractwise FA_perc_, MD_perc_,CBF_perc_, and ATT_perc_ were assessed using multivariate regression of sex, age and the age-sex interaction.

#### 2.2.2 Microstructure vs Perfusion

Two multivariate regression models were constructed to assess relationships between microstructure, perfusion, age, and sex in our sample. Both models were applied first to whole-WM MD_perc_ and FA_perc_ values, then to each tract:

● *Model 1. microstructure vs. perfusion across ages:* assesses the degree to which perfusion metrics account for the age-related variability in mean normalized tract MD and FA, with subject sex as a covariate of no interest. That is,

○ {FA_perc_, MD_perc_} = f(CBF_perc_+Sex)
○ {FA_perc_, MD_perc_} = f(ATT_perc_+Sex)
● *Model 2. microstructure vs. perfusion between sexes:* assess the degree to which perfusion metrics account for sex-related differences in normalized tract MD and FA, with subject age as a covariate of no interest. That is,

○ {FA_perc_, MD_perc_} = f(CBF_perc_+Age)
○ {FA_perc_, MD_perc_} = f(ATT_perc_+Age)
○ One set of models is produced for each sex.

The above analyses were conducted in R (version 4.1.1). False Detection Rate (FDR) multiple comparison corrections were conducted on a per-model basis for all regressions.

#### 2.2.3 Multimodal associations: Canonical correlation analysis (CCA)

Canonical correlation analysis (CCA) was conducted to examine the degree to which each of our variables of interest contributed to associations between microstructure and perfusion, respectively. CCA allows the creation of linear combination vectors of multiple related measures to examine the degree of contribution (or loading) of each constituent variable to the relationship between the tested vectors. Two column vectors were computed using our perfusion (*V*_1_) variables (CBF_perc_, ATT_perc_) and microstructural (*U*_1_) variables (AD_perc_, RD_perc_, FA_perc_) in Matlab. The linear combinations of *V* and *U* vectors which produced the highest correlation were then calculated to determine on a tractwise basis which variables loaded most strongly to correlated (between perfusion and microstructure) age differences by tract. Values more than three scaled median absolute deviations (*Difference of Median*, n.d.) from the median were excluded for each measure prior to CCA analysis. Split-half cross-validation was conducted through 10,000 permutations of random sampling of the male and female data sets. CCA was performed on each of the 10,000 randomly split samples, and mean and standard error of loadings associated with only statistically significant canonical correlations are plotted to confirm consistency of the CCA findings.

#### 2.2.4 Age-displaced analysis: cross-correlation function (CCF)

Cross-correlations functions (CCF) were calculated in Matlab (Mathworks, Natick, USA) to determine the presence of shifted correlations across measures for perfusion and microstructure. In the absence of longitudinal data, cross-correlation allows the examination of associations between variables at different points along cross-subject age-related trajectories, and the generation of hypotheses that could be tested longitudinally. In this work, a positive peak-correlation shift in variables A relative to variable B, indicates that variations in B are observable before they are observable in variable A (**Figure 2b**). Only variables demonstrating significant non-linear age effects on a tractwise basis were examined in this analysis to reduce ambiguity in CCF interpretation. A curvature threshold for age effects of 0.0005 was implemented, where curvature was computed using Matlab’s *curvature* function. CCFs were calculated based on the models of age effects for six paired tractwise microstructure/morphometry measures, along with cross-comparisons for AD_perc_/RD_perc_ vs FA_perc_ and CBF_perc_ vs ATT_perc_:

- CBF vs. Microstructure: CBF_perc_ vs AD_perc_, CBF_perc_ vs RD_perc_, CBF_perc_ vs FA_perc_
- ATT vs. Microstructure: ATT_perc_ vs AD_perc_, ATT_perc_ vs RD_perc_, ATT_perc_ vs FA_perc_
- AD_perc_ ^vs FA^_perc_
- RD_perc_ ^vs FA^_perc_
- CBF_perc_ vs ATT_perc_

**Figure 2:**
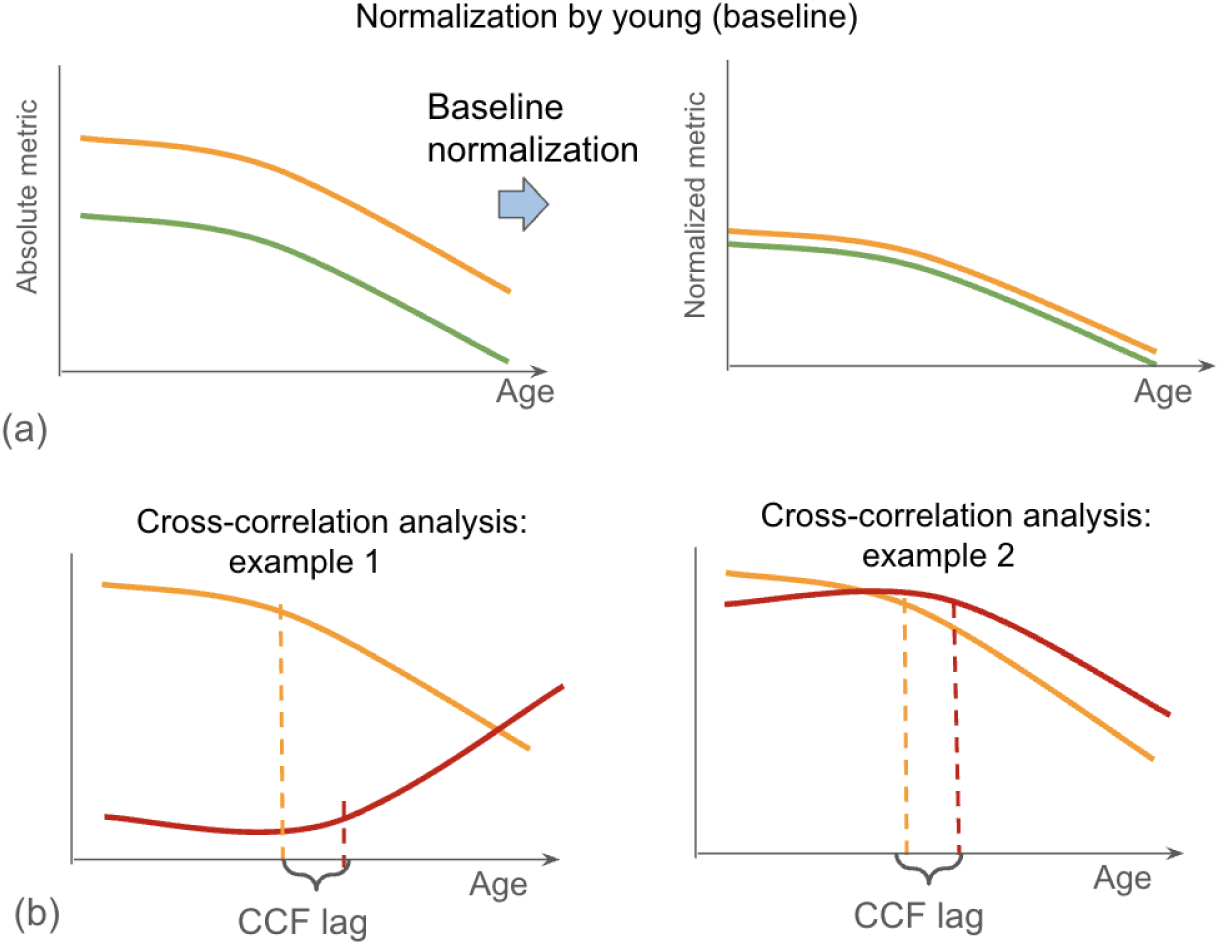
Illustration of a) relative effects of baseline normalization on study metrics to produce FA_perc_, MD_perc_, AD_perc_, RD_perc_, CBF_perc_, and ATT_perc_, values, as well as b) interpretation of peak offset in cross correlation analysis.

## 3 RESULTS

### 3.1 Age differences

Significant linear and quadratic relationships between age and CBF_perc_ were identified in all tracts but UNC, with quadratic regressions providing a better fit as indicated by higher *R^2^* in all tracts with the exception of Fminor and CCG. Significant associations were also identified between age and ATT_perc_ in Fmajor, ILF, and UNC, along with significant linear effects in Fminor, ATR, CCG, CST, SLFP, SLFT, and UNC, with quadratic regression providing a better fit in UNC. Significant linear and quadratic relationships between age and MD_perc_ were identified in all tracts of interest, with quadratic regressions providing a better fit in all tracts. Lastly, significant quadratic associations between age and FA_perc_ were found in Fminor, CCG, and ILF, with significant linear associations in Fmajor, Fminor, ATR, CAB, CCG, ILF, SLFP. Quadratic regression provided a better fit for the effects of age on FA in Fminor and CCG (**Figure 3**).

To further verify the viability of ATT estimation in the WM, we found that across subjects, both CBF and ATT values in the whole WM were linearly associated with those across the whole GM (**Supplementary** Figure 1). Linear regression of GM CBF/ATT as a function of WM CBF/ATT returned a strong association between tissue types: slope = 0.13 (WM CBF vs. GM CBF), intercept = 2.4, *F*(1, 498) = 633.4, *p* < .001, *R*^2^_Adjusted_ = .56; slope = 0.26 (WM ATT vs. GM ATT), intercept = 1.3, *F*(1, 498) = 256.5, *p* < .001, *R*^2^_Adjusted_ = .34. Notably, WM ATT was on average 1.3 s longer than GM ATT, which is consistent with the range published in the literature (Juttukonda et al., 2021; van Osch et al., 2009).

**Figure 3:**
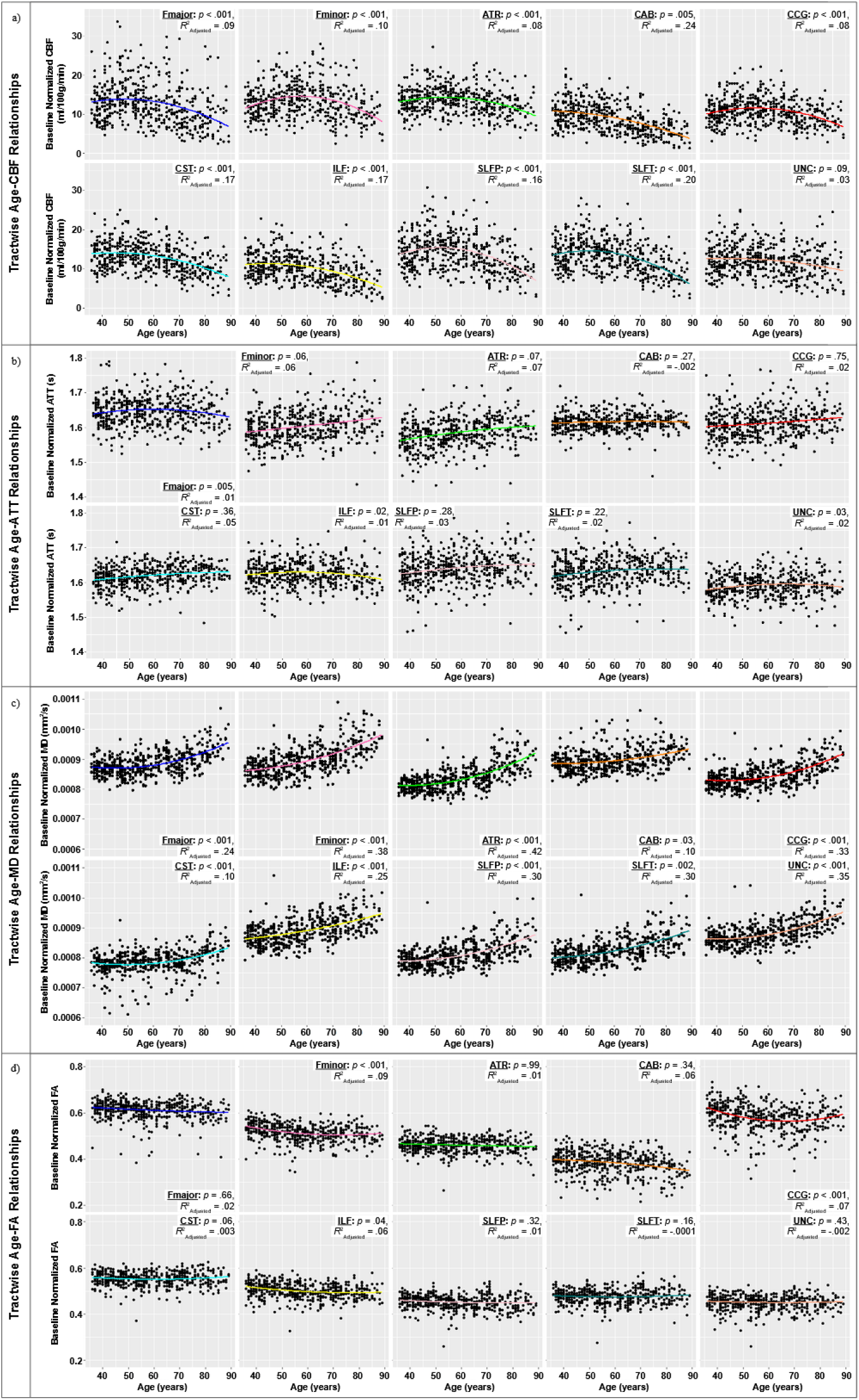
Tractwise quadratic regression of age effects on baseline normalized CBF, ATT, MD, and FA, with each dot representing one subject. The p-values and effect sizes (R2adjusted) from the quadratic fit are noted for each tract.

**Figure 4:**
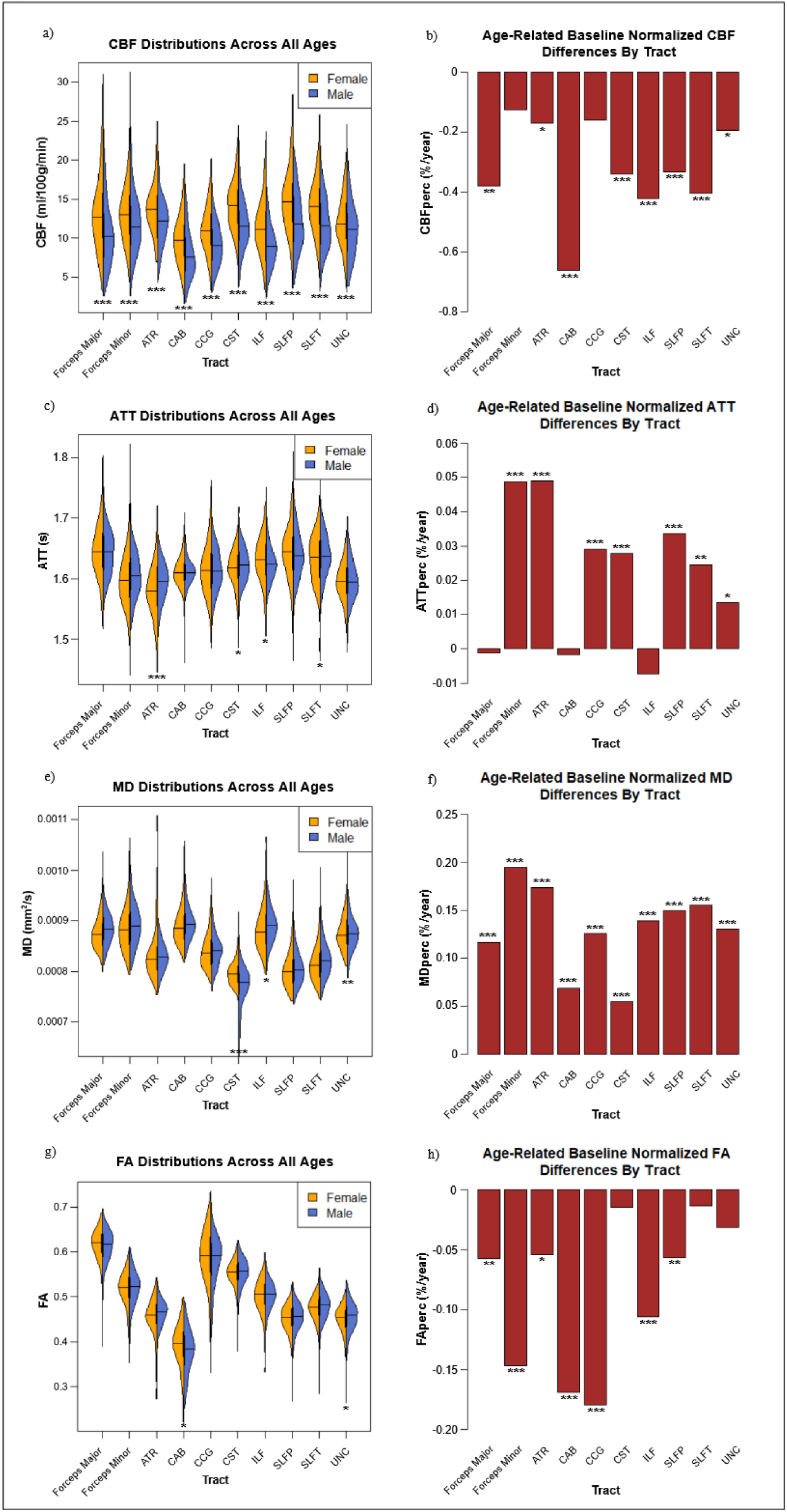
Raw distributions separated by sex (a,c.e,g) and tractwise age effects (b,d f.h) for CBF_perc_, ATT_perc_, MD_perc_, and FA_perc_ respectively. Significant effects by sex and age are denoted with asterisks (*:p<.05, **p<.01, **:p<.001)

### 3.2 Sex differences

Prior to baseline normalization, higher raw MD was found in female subjects in the CST, and in male subjects in the ILF and UNC. FA differences were similarly mixed where significant, with female subjects showing higher FA in the CAB, and male subjects showing higher raw FA in the UNC. More consistent sex differences were found in CBF, with female subjects showing higher raw CBF in all tested tracts, corroborating previous research on sex differences in cortical perfusion (Y. Liu et al., 2012; Tosun et al., 2010). However, raw ATT values again demonstrated tract dependence in sex differences, with longer raw ATT in female subjects in the ILF, and longer raw ATT in male subjects in the ATR, CST, and SLFT (**Figure 4**).

Following baseline normalization, female subjects showed significantly higher MD_perc_ in the Fmajor, ATR, CST, ILF, SLFP, and SLFT in regression analysis. That is, these tracts may demonstrate more microstructural decline with advancing age than the others. While female subjects also demonstrated higher FA_perc_ in the Fmajor, male subjects showed higher FA_perc_ in the Fminor, CCG, SLFP, and SLFT. Female subjects demonstrated higher CBF_perc_ in the CAB, while male subjects showed higher CBF_perc_ in the ATR. Female subjects showed higher ATT_perc_ in the SLFP and SLFT, while male subjects showed higher ATT_perc_ in the ILF.

Significant age-sex interactions were identified for MD_perc_ in the Fminor (*F*(2, 489) = 70.8, *p* = .007, *R*^2^_Adjusted_ = .30), ATR (*F*(2, 493) = 76.9, *p* = .009, *R*^2^_Adjusted_ = .29), CAB (*F*(2, 486) = 11.5, *p* = .046, *R*^2^_Adjusted_ = .06), and UNC (*F*(2, 491) = 47.3, *p* = .015, *R*^2^_Adjusted_ = .22), such that in all instances, female subjects exhibited greater MD increases in aging. Interactions of age effects by sex were also found in FA_perc_ in the ATR (*F*(2, 493) = 4.37, *p* = .013, *R*^2^_Adjusted_ = .02), SLFP (*F*(2, 493) = 6.03, *p* = .009, *R*^2^_Adjusted_ = .03), and SLFT (*F*(2, 491) = 5.60, *p* = .001, *R*^2^_Adjusted_ = .03), with greater FA_perc_ decreases in aging for female subjects. Interactions of age by sex for CBF differences were only identified in the CCG (*F*(2, 491) = 2.49, *p* = .042, *R*^2^_Adjusted_ = .009), such that CBF_perc_ declines were greater in female subjects. No interactions by subject sex were found in age-related ATT_perc_ differences.

### 3.3 Microstructure-perfusion relationships

#### 3.3.1 Regression analyses

*Model 1 (perfusion vs microstructure):* Significant relationships were identified between CBF_perc_ and MD_perc_ (*F*(2, 497) = 5.36, *p* = .005, *R*^2^_Adjusted_ = .02, slope = -0.02), as well as between ATT_perc_ and MD_perc_ (*F*(2, 497) = 4.07, *p* = .02, *R*^2^_Adjusted_ = .01, slope = 0.23). No significant whole-WM relationships were found between FA_perc_ and WM perfusion (**Supplementary Table 1a**).

Significant negative relationships were found between tractwise CBF_perc_ and MD_perc_ in the ILF and SLFT. Significant positive relationships between ATT_perc_ and MD_perc_ were identified in the Fminor, ATR, SLFP, and SLFT. A single significant positive association was found between CBF_perc_ and FA_perc_ in the CCG. A significant negative relationship between ATT_perc_ and FA_perc_ was identified in the ATR, along with a positive relationship in the CAB Tract-wise results are summarized in **Supplementary Table 1b**.

*Model 2 (sex differences in perfusion-microstructure relationships):* No significant relationships were identified between whole-WM microstructure and perfusion in Model 2 (**Supplementary Table 2a**). After dividing the sample by sex, significant positive associations were found between CBF_perc_ and FA_perc_ in the CCG in both female and male subjects, as well as in the ATR in female subjects alone. A single negative association between CBF_perc_ and FA_perc_ was found in the CST in male subjects. Female subjects also demonstrated both a negative association between ATT_perc_ and FA_perc_ in the ATR, and a positive association between ATT_perc_ and FA_perc_ in the CAB. No associations were identified between perfusion and MD after dividing the sample by sex. Tract-wise results are summarized in **Supplementary Table 2b.**

#### 3.3.2 Multimodal associations based on CCA

The CCA identified a common pattern of association between microstructure and perfusion loadings in male and female subjects in several tracts (**Figure 5**). Loadings are listed for significant canonical correlations (p <0.05) using a loading threshold of 0.3. In female subjects, CBF_perc_ contributed significantly in six tracts (ATR, CCG, CST, ILF, SLFP and UNC), while ATT_perc_ did not contribute significantly to perfusion-microstructure associations in any tract. Across tracts, AD_perc_ and RD_perc_ were not found to load significantly, while FA_perc_ loaded significantly in four tracts (Fminor, ATR, CCG and UNC). Three tracts demonstrated significant loading by both CBF_perc_ and FA_perc_ (ATR, CCG, UNC) in females.

In male subjects, CBF_perc_ also contributed significantly to the perfusion-microstructure relationship in six tracts (ATR, CCG, CST, ILF, SLFP, SLFT), while FA_perc_ co-contributed significantly in only one tract (CCG). The CCG was also the only tract displaying significant loading by both CBF_perc_ and FA_perc_ for males. No other measures demonstrated significant loading in male subjects. In both sexes, FA_perc_ was the only microstructural parameter to contribute to these relationships, while CBF_perc_ showed the only significant perfusion loading.

**Figure 5:**
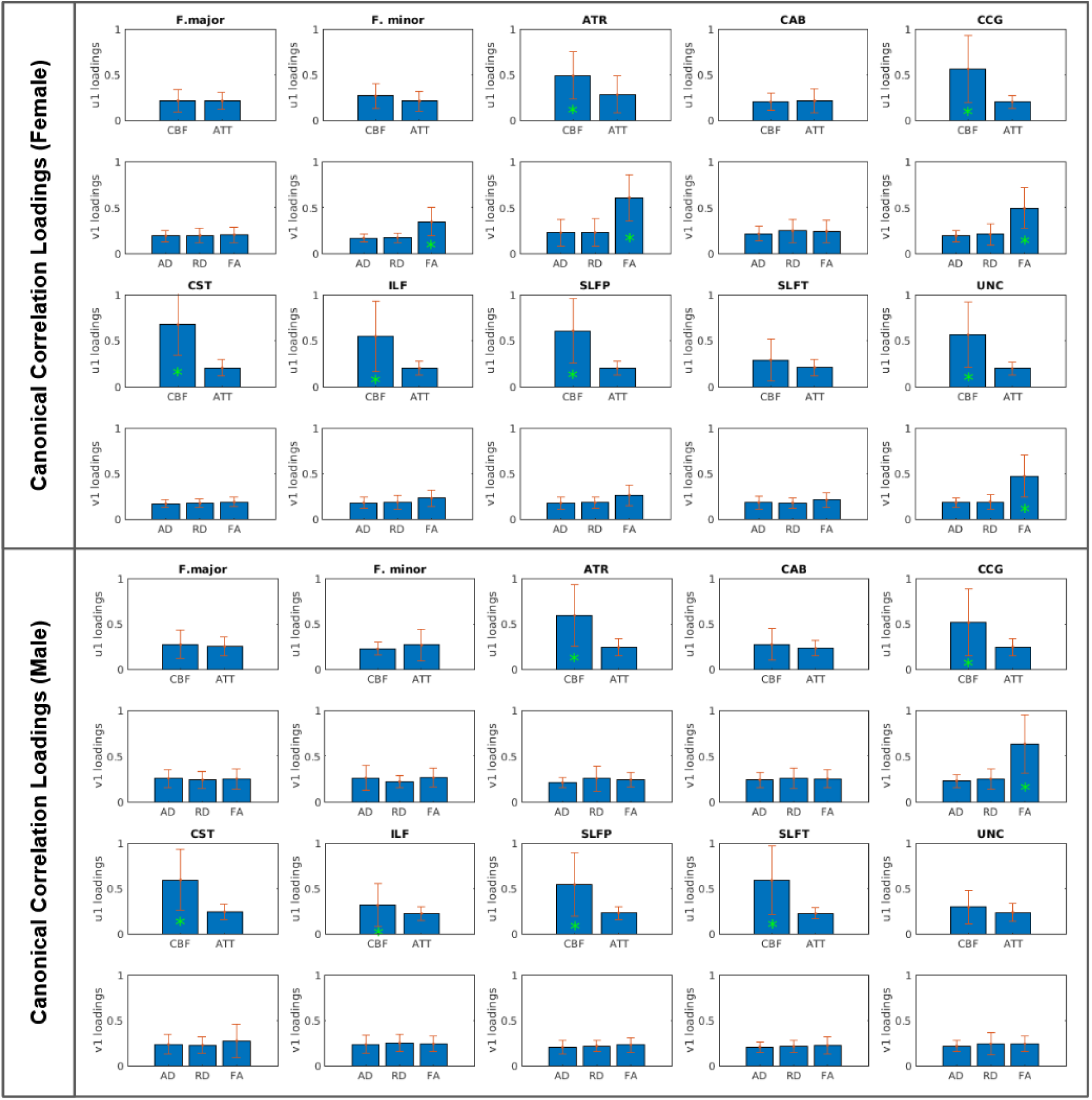
Tract-wise contributions by all perfusion (CBF_perc_, ATT_perc_ ) and microstructural AD_perc_, RD_perc_, FA_perc_) measures to categorical perfusion-microstructural relationships in aging. U1 and V1 refer to latent variables related 10 perfusion and microstructure, respectively, Green asterisks denote the loading threshold of 0.3.

**Figure 6:**
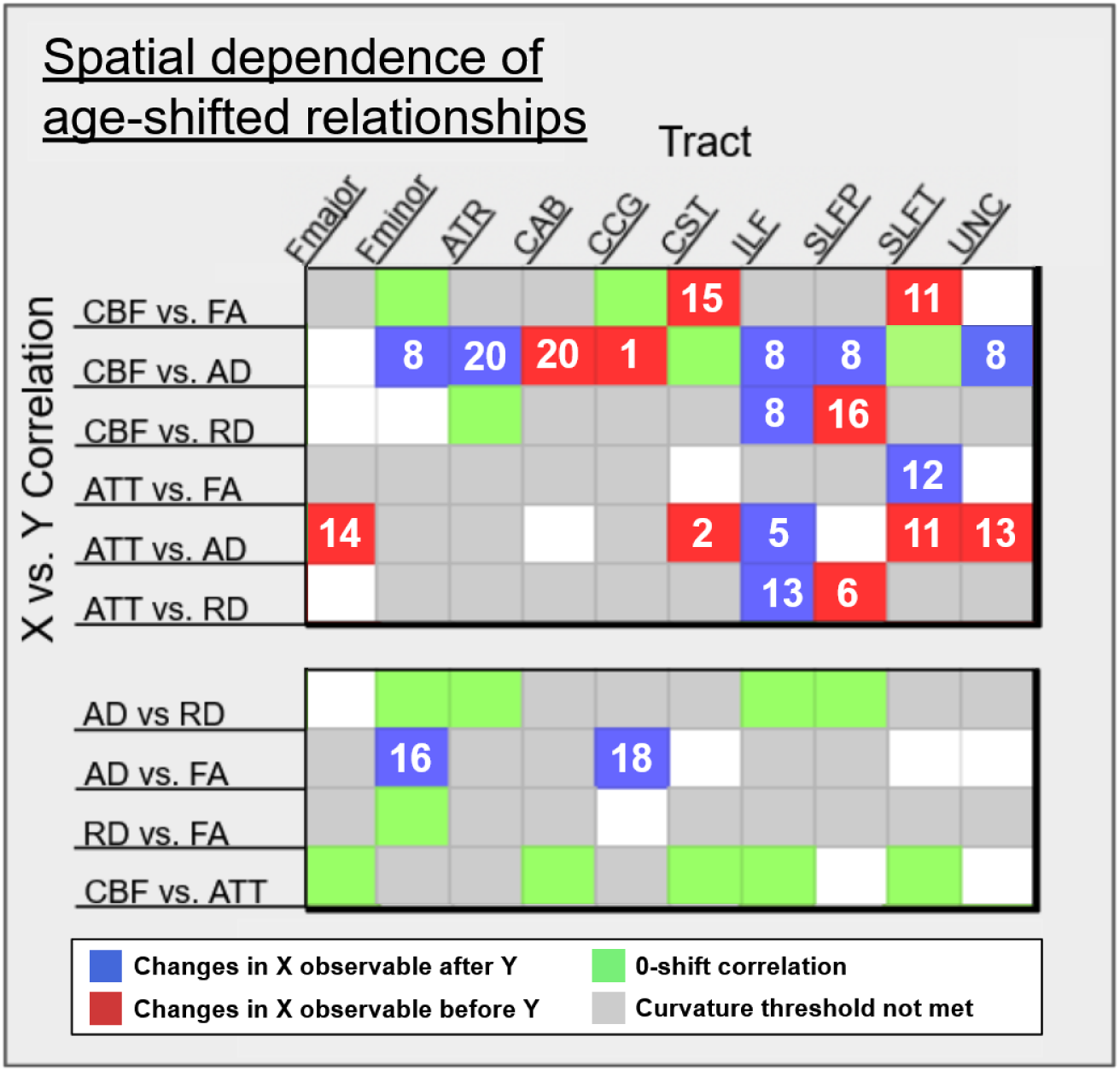
Tractwise cross-correlation results demonstrating significant peak correlations (p < .05) for pairs of perfusion and microstructural variables at positive, negative and zero correlational shift. While cells indicate no significant association between measures. Length of peak-offset in years (approx.) is included in relevant cells.

#### 3.3.3 CCF results of age-displacement analysis

Correlating perfusion and microstructure measures across age shifted timeseries identified a broad ordered trend in earliest detectable ages of change of ATT_perc_, followed by AD_perc_, then CBF_perc_ and the remaining microstructural measures, in addition to considerable tractwise differences.

Significant correlations at zero peak-correlational shift were observed between CBF_perc_ and FA_perc_ in Fminor and CCG, between CBF_perc_ and AD_perc_ in CST and SLFT, and between CBF_perc_ and RD_perc_ in the CAB. Significant associations at a positive peak-correlation shift were found in CBF_perc_ relative to FA_perc_ in CST and SLFT; between CBF_perc_ and AD_perc_ in CAB, and CCG; and between CBF_perc_ and RD_perc_ in the SLFP, such that CBF_perc_ changes were observable at earlier ages than FA_perc_. In other words, in these tracts, variations in CBF_perc_ were observable before those in FA_perc_ and AD_perc_. Similarly, significant correlations at positive shift between ATT_perc_ and AD_perc_ were found in Fmajor, CST, SLFT, and UNC; and between ATT_perc_ and RD_perc_ in the SLFP, such that variations in ATT_perc_ were observable before those in AD_perc_ or RD_perc_. Significant correlations at negative shift were found between CBF_perc_ and AD_perc_ in Fminor, ATR, ILF, SLFP, and UNC, as well as between CBF_perc_ and RD_perc_ in the ILF alone, such that variations in CBF_perc_ were observable after those in AD_perc_ or RD_perc_. Significant correlations at negative shift were also found between ATT_perc_ and FA_perc_ in the SLFT; between ATT_perc_ and AD_perc_ in Fmajor, CST, SLFT, and UNC; and between ATT_perc_ and RD_perc_ in the ILF_perc_ alone such that changes in ATT_perc_ were observable at later ages than changes in FA_perc_, AD_perc_, or RD_perc_ respectively (**Figure 6**).

Comparisons within perfusion and microstructure measures respectively found significant correlations at zero shift between AD_perc_ and RD_perc_ in Fminor, ATR, ILF, and SLFP; as well as between RD_perc_ and FA_perc_ in the Fminor. In perfusion measures, significant zero-shift correlations were found between CBF_perc_ and ATT_perc_ in Fmajor, CAB, CST ILF, and SLFT. Additionally, two significant correlations at negative peak shift were identified between AD_perc_ and FA_perc_ in Fminor and CCG, such that changes in AD_perc_ were observable at earlier ages than changes in FA_perc_.

## 4 DISCUSSION

We initially hypothesized that (1) tractwise microstructural and perfusion declines would not progress at the same rate in aging due to (2) earlier detectable declines in WM perfusion than microstructural integrity. Our findings support Hypothesis 1 and in part Hypothesis 2, but also highlight the fact that the patterns of decline were much more varied by tract, sex and by measure than initially stated in Hypothesis 2.

MD was largely found to increase quadratically with age, while FA declined linearly in the majority of tracts. While this finding is in broad agreement with the aging literature (de Groot et al., 2016; Giorgio et al., 2010; Slater et al., 2019), FA was not uniformly associated with age across tracts, indicating variability in patterns or rates of WM decline. Across sexes, our microstructural results provide partial support for fronto-temporal tracts declining at a higher rate than those in the posterior WM (**Figure 4**). We also found significant and consistent CBF and ATT associations with age. Specifically, CBF declines quadratically with advancing age in the vast majority of WM tracts, somewhat mirroring the age-associated increases in tractwise MD (**Figure 3**), with the quadratic behaviour consistent with previous findings by Juttukonda et al. in the same cohort (Juttukonda et al., 2021). CBF appears to demonstrate a peak in middle-age, with the peak location varying by tract. Moreover, ATT demonstrates a robust linear increase with advancing age, consistent with previous reports (Mutsaerts et al., 2015).

Importantly, we found a tendency for ATT to lead microstructural metrics in age-related variations. Nonetheless, this effect is tract-dependent, which may be explained by the tract-dependent cellular and vascular developmental processes.

### 4.1.1 Tract-based approach

With regards to vascular development, vessels in frontal and cerebrospinal tracts develop in the embryo prior to those in the parietal and posterior brain, supporting the development of their respective WM regions (Smirnov et al., 2021). This raises vascularization differences as a potential source of differences in rates and patterns of WM declines. These lines of evidence encourage a tract-wise determination of perfusion-microstructural associations, whereby those tracts or regions that demonstrate greater robustness to age-related microstructural declines may be distinguished by vascular factors that are potentially related to the manner in which blood vessels developed. Unlike the cortical surface, which is fed by larger penetrating vessels, blood supply to the WM is fed primarily by smaller vessels entering from either the cortical surface or the midbrain (Nonaka et al., 2003; Smirnov et al., 2021). This results in both (1) considerable variation in proximity to major vessels and number of blood flow sources and (2) the likelihood that vascular disease and age-related deterioration that differentially affects small vessels has an earlier impact on the WM than on the cortical surface (Akashi et al., 2017; Hase et al., 2018). Tractwise heterogeneity in blood vessel development may then underlie susceptibility to age-related changes in perfusion and regional ability to support microstructural integrity in adulthood.

Thus, we took a tract-wise approach that complements recent work based on a voxel-wise approach (Damestani et al., 2024). While prior findings have indicated regional gains in WM CBF with aging (Alisch et al., 2021), we found significant trackwise CBF declines. The highest age effects on CBF appear to manifest in Fmajor, CAB, ILF and SLFT, suggesting higher rates of vascular decline in the temporo-posterior WM than more anterior-superior tracts such as the Fminor or CCG which demonstrate little to no CBF declines with age. Thus, vascular age effects do not appear to follow the current retrogenesis models that predict earlier or more rapid age declines in anterior regions (Brickman et al., 2012; Slater et al., 2019; Vik et al., 2015). However, patterns of age-related ATT increases across our tracts do not follow this pattern. Despite showing some of the greatest CBF declines, Fmajor, CAB, and ILF tracts exhibited no significant ATT increases with age. This pattern is in line with the voxelwise findings by Damestani et al. (Damestani et al., 2024), and suggests that ATT and CBF cannot be considered interchangeable measures of microvascular decline, and instead reflect different signatures or stages of microvascular aging that will require targeted analysis to clarify.

### 4.1.2 Understanding different types of WM-perfusion associations through multimodal integration

As revealed by the CCA, the interrelation between perfusion and microstructural integrity varied by tract. This highlights the importance of investigating the regional heterogeneity in the interplay between vascular and structural declines. Moreover, rather than a smooth shared trajectory of decline, our evidence indicates the presence of different combinations of age effects: (i) significant loadings by CBF_perc_ alone; (ii) significant loading by FA_perc_ alone; (iii) significant loadings by CBF_perc_ and FA_perc_ (**Figure 5**). In most cases, CBF_perc_ and FA_perc_ loaded in the same direction, consistent with the global relationship shown by Damestani et al. (Damestani et al., 2024). However, several tracts (Fmajor, CAB and SLFT in females, Fmajor, Fminor, CAB and UNC in males) did not show significant loading contributions by either microstructural or perfusion metrics. An interesting follow-up question is, given the obvious differences in variables such as perfusion and axonal diameter across these tracts, what mechanism(s) may be driving the heterogeneity in the perfusion-microstructural associations?

Notwithstanding, the understanding of the WM aging process appears to be generalizable across all WM. For example, using the CST as a point of reference for a tract presumed to survive relatively robustly into later age (Michielse et al., 2010), we see that the CST exhibits the lowest MD increases and no significant differences in FA with age, despite sizable CBF_perc_ and ATT_perc_ age effects. As axon diameter comparisons across the WM have observed relatively high mean diameter in CST fibers (Fan et al., 2020; Liewald et al., 2014), fiber diameter in healthy WM may be a predictor for resilience to perfusion declines. Additionally, tracts with different mean axon diameters (Fan et al., 2020; Gast et al., 2023) are accompanied by differing metabolic demands (Harris & Attwell, 2012). One possibility for this is that the metabolic cost of maintaining myelin itself scales with fiber diameter, so thicker fibers, such as those of the CST may be less metabolically sensitive than those with thinner fibres, such as the UNC. Nonetheless, the driver of sex-related differences in these vascular-microstructural differences remain unclear.

### 4.1.3 Sex differences in perfusion-microstructural covariations

In addition to perfusion-microstructure associations differing by tract, they also varied considerably between sexes (**Figure 5**). First, female subjects have poorer WM integrity as a group, as shown by the higher group-averaged MD_perc_ and lower FA_perc_ values (**Figure 7**), and as shown previously (Inano et al., 2011; Intzandt et al., 2023; Lawrence et al., 2021). Females also tend to have higher CBF_perc_ (consistent with previous findings (Alisch et al., 2021)) and shorter ATT_perc_ (in anterior tracts) compared to males (**Figure 4**). Specific to multi-delay pCASL-based findings, these are similar to observations by Damestani et al. (Damestani et al., 2024; Juttukonda et al., 2021), although our values are normalized by the youngest group and separately for each sex, implying that in addition to sex differences in regional baseline (young) values, the regional age-related variations are also sex dependent. Moreover, female subjects demonstrated a greater number of tracts with significant relationships between WM microstructure and perfusion. This finding is consistent with previous reports of more rapid WM aging in females (Intzandt et al., 2023; Patel et al., 2023). We speculate that a likely cause of these differences in WM aging by sex are the effects of menopause on brain health more broadly. Menopause has previously been found to affect structural, connectivity, and metabolic measures in the brain (Mosconi et al., 2021).

Notably, cognitive impacts appear to stabilize post-menopause, though this is speculated to be the result of compensatory effects, rather than improvement in structural integrity. The sex differences with age observed in this study likely capture structural and microvascular impacts of peri- and postmenopause on the WM, though direct associations between these declines and cognitive performance will require additional investigation. Should microvascular declines be associated with cognitive performance similarly to the microstructural associations identified in this study, targeting vascular declines may benefit the study of interventions in the impacts of menopause on the brain in addition to the aging brain more generally. The degree to which hormone supplementation therapies may mitigate these effects, and indeed the mechanisms of estrogen’s effects on WM health more generally, will require further investigation.

**Figure 7:**
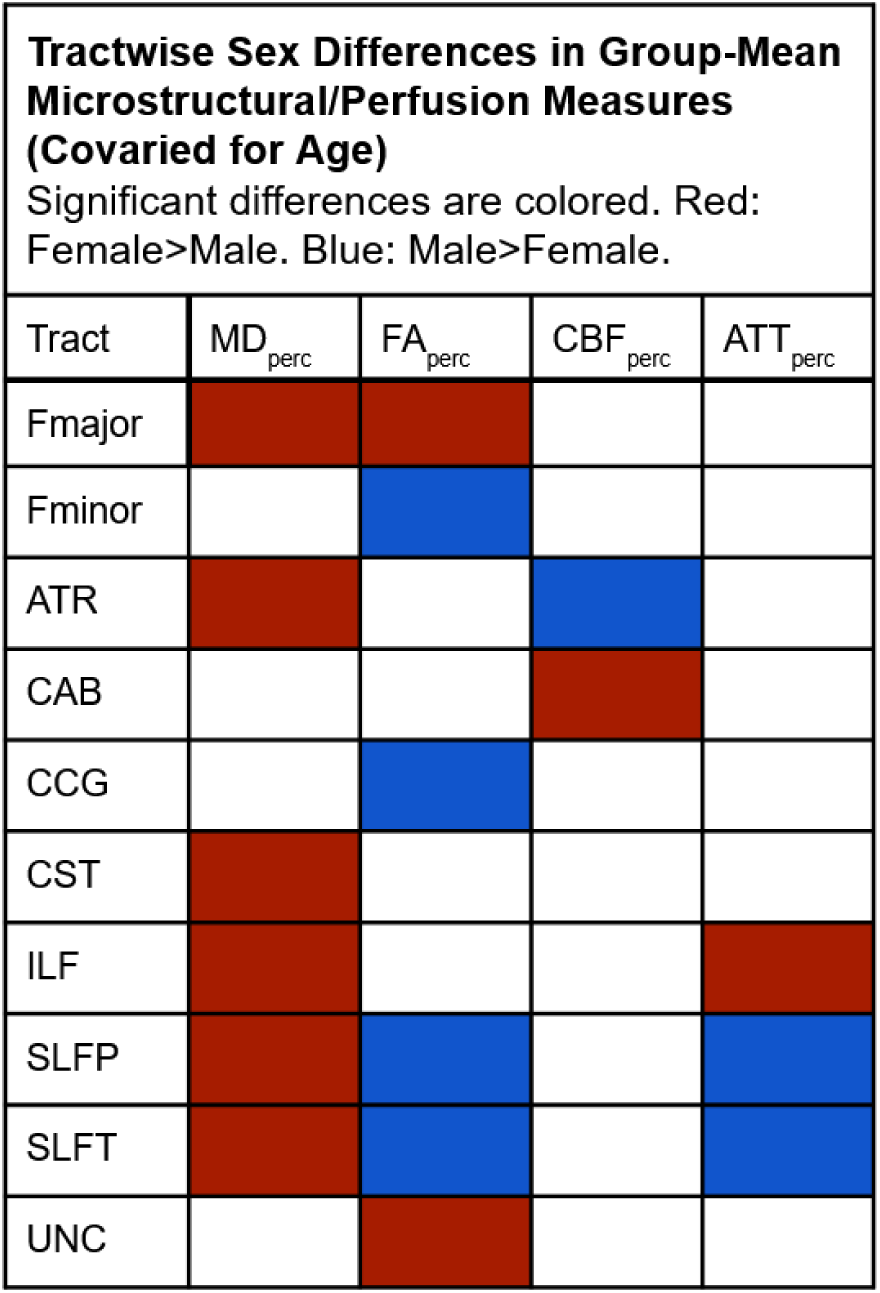
Tractwise sex differences in baseline normalized CBF, ATT, MD, and FA.

### 4.1.4 Temporal relationship in perfusion-microstructural variations in aging

Vascular declines with age have been consistently demonstrated in the WM as part of general systemic vascular decline (Kohn et al., 2015; Y. Li et al., 2018; Tomoto et al., 2023). Increasing arterial stiffness due to the progressive decay of elastin in normal aging results in more fragile blood vessels (Kohn et al., 2015; Tomoto et al., 2023). Even in intact arterial pathways, changes in vessel morphometry result in more tortuous micro- and mesovasculature with reduced blood flow (Y. Li et al., 2018). As a result, diminished perfusion into WM tissues is expected in healthy aging, and are suggested to underlie microstructural changes (Bagi et al., 2018; Chen et al., 2013). The results (**Figure 6**) of our cross-correlation analysis are key to the vascular hypothesis of WM aging, and indicate that age-related deterioration may be detectable via ATT prior to most microstructural measures in certain tracts (particularly in the CST, SLFT, SLFP and UNC). Following this, AD_perc_ is the microstructural measure displaying detectable variations at the youngest ages. CBF_perc_ declines, in contrast, were detected at older ages than AD_perc_. These lead us to hypothesize that ATT may be an earlier marker of WM decline than microstructure and WM CBF, to be confirmed in a longitudinal design. The superior sensitivity of vascular timing information to pathology would be consistent with recent reports on the unique sensitivity of vascular-response latency (Holmes et al., 2020; D. Kim et al., 2021; McKetton et al., 2019; Nanayakkara et al., 2023). In fact, ATT has previously been found to exhibit lower variability than CBF (Mutsaerts et al., 2015), which may underlie the finding that ATT, not CBF, displayed a detectable earlier variation in aging. Of the regions showing a leading variation in ATT_perc_ in the CCF analysis, the age associations with ATT_perc_ or any of the other parameters were not all particularly high or low, arguing against a detectability bias due to age effect size (**Figure 4**). The only tract in which ATT_perc_ variations lag those in microstructure is the ILF. As prior research has demonstrated an association between reduced perfusion in the cortex and the development of WM hyperintensities in related WM regions (Mayer et al., 2021), it is intuitive that changes in perfusion in the WM precede declines in WM integrity and ultimately the breakdown of WM tissue. This process likely results from the progressive inability of supporting glial cells to maintain myelin in WM fibers as perfusion (CBF) declines (Kobayashi et al., 2002; Kohama et al., 2012). Those axons with reduced or malformed myelin sheaths are then more vulnerable to age-related culling of vulnerable WM fibers, and associated WM regions at higher risk of pathological declines (Kohama et al., 2012; Tang et al., 1997). Related to cerebrovascular mapping, vascular response latency has been demonstrated as a more sensitive marker of pathology than response amplitude (Holmes et al., 2020; D. Kim et al., 2021; McKetton et al., 2019; Nanayakkara et al., 2023). Additionally, arterial-transit time (ATT) has been found to increase with advancing age (Mutsaerts et al., 2015). Moreover, arterial-transit time is reportedly independently related to white-matter lesions relative to CBF (Zhang et al., 2018). Thus, it is also interesting to determine whether CBF or ATT is an earlier marker of possible WM degeneration.

However, the ILF appears to be an exception in our results, in that all perfusion metrics lag microstructural ones. The ILF is characterized by a particularly low ATT_perc_, but in general, we noted that the strength of the ATT_perc_ age effects is not associated with group-mean ATT_perc_ values, so the differences in the ATT’s temporal evolutions across tracts remains unexplained. It is interesting to note that the ILF is the one most connected to the temporal cortex and temporal pole, among the earliest regions to deteriorate in aging (Cox et al., 2021), and a region with accelerated deterioration in cases of mild cognitive impairment and associated with early stages of Alzheimer’s disease (X. Li et al., 2020; Wu et al., 2021).

Thus, it is possible that the temporal brain tissue is differently susceptible to the aging process. More generally, glial function has been reported to deteriorate in aging independently of vascular factors (Angelova & Brown, 2019; Hanslik et al., 2021; Lau et al., 2023; Rawji et al., 2023). Clarifying peak-correlation shifted relationships in those tracts exhibiting linear declines will require either a wider age sample able to capture younger peak ATT values, or analysis of longitudinal data. A more comprehensive understanding of these differences would require accounting for physiological and developmental variation between tracts (REF paper 3). Nonetheless, this is the first time that WM perfusion has been directly demonstrated to vary at an earlier age than WM microstructure. We theorize that CBF and ATT reflect different aspects of vascular aging. However, vascular decline is fastest in the temporo-posterior tracts, while WM microstructural decline is fastest in the fronto-temporal tracts, suggesting vascular decline is not the sole contributor of WM decline. Future investigations should additionally consider the role of the anterior-posterior retrogenesis hypotheses (Hoagey et al., 2019; Raz, 2000; Slater et al., 2019).

### 4.1.5 Limitations

Our CCF analysis depends on nonlinearity in the age-associations of its inputs. While many tracts demonstrated significant quadratic declines, the heterogeneity of significant quadratic versus linear age effects restricted the tracts in which cross-measure tractwise comparisons could be conducted. Moreover, a notable limitation to these findings is that, while CBF_perc_ declined quadratically in all tracts but the UNC, the predominantly linear declines in ATT_perc_ restricted the extent of possible timing comparisons. In any case, validating the temporal relationships and causal relationships will require the inclusion of longitudinal data.

One interesting possibility of WM perfusion findings in general is that different tracts, which may have become vascularized to different extents and at different developmental stages (Smirnov et al., 2021), may be differentially susceptible to degeneration. As cerebral microbleeds (CMB) are present in an estimated 23.5% of the healthy aging population (Litak et al., 2020; Pasi et al., 2017), rising from 17.8% in ages 60-69 to 38.3% over age 80 (Ding et al., 2015; Jeerakathil et al., 2004), there is little doubt that a degree of cerebral small vessel disease (CSVD) is present in this sample. The prevalence of these conditions in typical aging precludes their removal from a representative sample. CMB have been associated with a loss in general microstructural integrity consistent with the microstructure-microvascular associations indicated by this study (J.-Y. Liu et al., 2020), however, the local impact of possible lacunes on our sample is limited by TRACULA’s deterministic probability tractography approach with anatomical priors (Maffei et al., 2021; Yendiki et al., 2011). Thus, for all practical purposes, tracts will not be interrupted even in the presence of CSVD. This was demonstrated in a recent paper by Maffei et al. (Maffei et al., 2023). Regardless, in future studies, we will also probe the effect of thresholding on segmentation and the differences between those with and without heavy CMB burden.

While evidence indicates the presence of notable impacts of menopause on WM integrity (Mosconi et al., 2021), self-report menopause status included in this data set was not sufficiently accurate to include in statistical analyses. Inclusion of more precise reporting of pre-, peri-, or post-menopausal status would be of considerable interest to furthering investigation of sex differences in WM aging. More generally, this study aimed to lay the groundwork for multimodal integration of aging metrics in the white matter and chose to address widely applied microstructural measures such as MD and FA. This introduces a number of limitations that may be resolved in more advanced diffusion modeling. This study also addressed only ten major WM tracts. These were chosen to minimize tract overlap/partial-voluming between segmentations, however, additional comparisons may be possible in more recent segmentations (Maffei et al., 2021). Finally, while we have identified novel age-related findings using our paradigm, future exploration using longitudinal data would complement the cross-sectional nature of the HCP-A dataset used in this study.

### 4.2 Conclusions

Relationships between age, perfusion, and microstructural integrity assessed in this study depict highly tract-specific, as well as sex-specific, variation across our sample. The lack of a unified set of trends between perfusion and microstructural declines across tracts supports our hypothesis that WM perfusion and microstructure do not decline at the same rate in aging. Additionally, our secondary hypothesis that WM perfusion differences would be detectable earlier than differences in microstructural integrity, found support in that tractwise age effects on ATT_perc_ appear at earlier ages than multiple microstructural measures. Moreover, the ubiquity of CBF_perc_ as the chief driver of significant canonical perfusion-microstructure relationships demonstrates that CBF and ATT reflect different aspects of WM microvascular aging and should be investigated in more depth. To our knowledge, this study is the first demonstration of tractwise age effects in WM CBF and ATT, and these findings will be highly valuable to the development of a more complete model for mechanisms of brain aging.

## Acknowledgments

We are grateful for the financial support from the Vascular Training (VAST) Platform (TDR), the Canadian Institutes of Health Research (CIHR) (#PJT169688) and the Canada Research Chairs (CRC) program (JJC).

## Data Availability

Data analyzed in this study was drawn from the Human Connectome Project in Aging (HCP-A) (Bookheimer et al., 2019). All included HCP-A datasets were made available under appropriate data usage agreements and can be accessed via the NIMH Data Archive (https://nda.nih.gov/general-query. html?q=query=featured-datasets:Human%20Connectome%20Projects%20(HCP)).

## Author Contributions

**T. D. Robinson:** Conceptualization, Methodology, Software, Formal Analysis, Data Curation, Writing - Original Draft, Visualization. **Y. L. Sun:** Software, Data Curation, Writing - Review & Editing. **P. T. H. Chang:** Software, Data Curation. **C. J. Gauthier:** Writing - Review & Editing. **J. J. Chen:** Conceptualization, Methodology, Formal Analysis, Writing - Review & Editing, Supervision, Project administration, Funding acquisition.

## Declarations

### Consent for Publication

All authors have consented to the publication of this study and its findings as an original research article in the journal GeroScience.

### Ethics Statement

All subject data included in this study was drawn from the Human Connectome Project In Aging

(HCP-A) (Bookheimer et al., 2019). All subjects provided informed consent following assessment of their capability to provide said consent and these data were made available for use under appropriate data usage agreements.

## Supplementary Materials

**Supplementary Figure 1:**
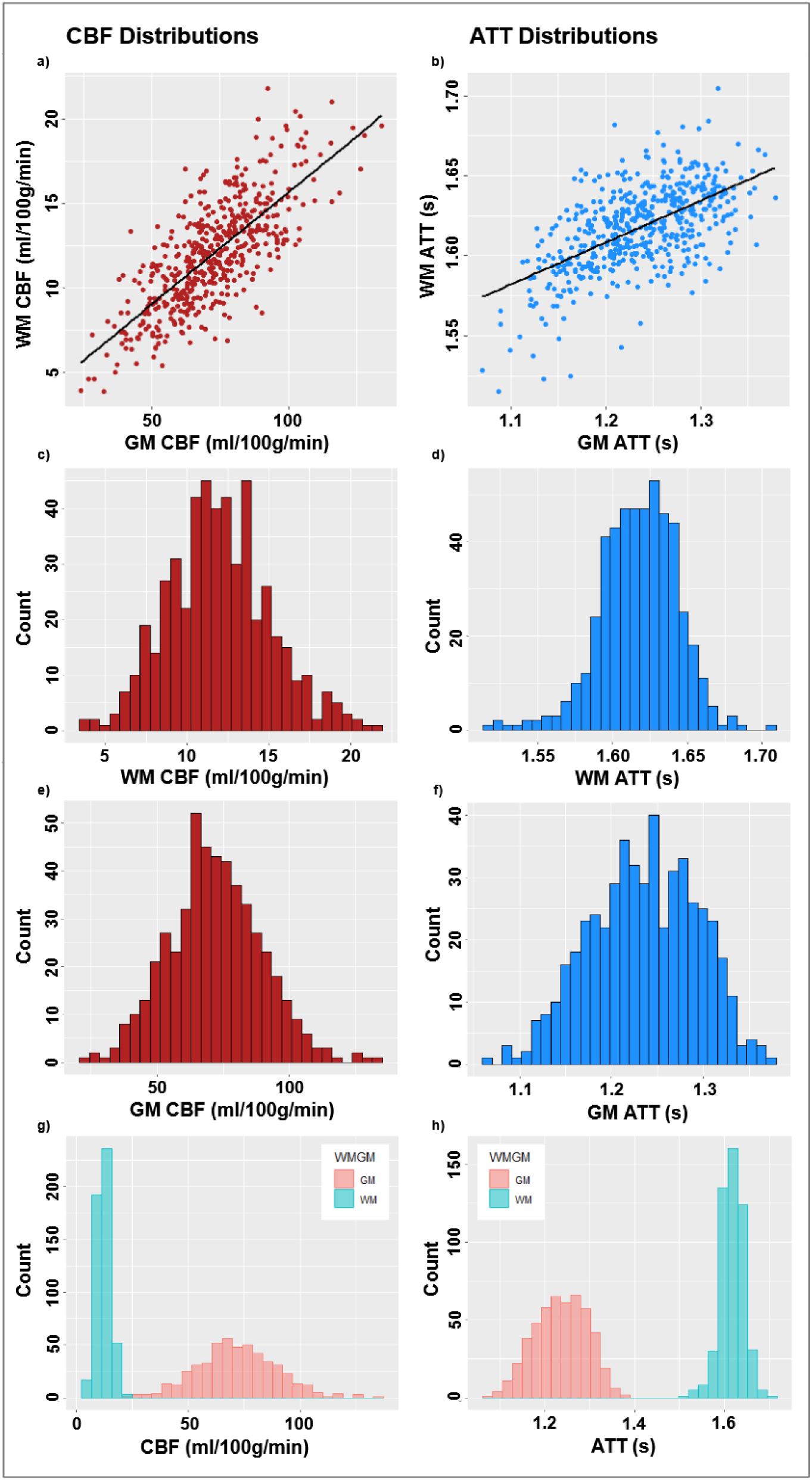
CBF and ATT distributions displayed between WM and GM values per subject (a, b), as separate counts for WM (c, d) and GM (e, f) across WM/GM specific ranges, and combined counts for both WM and GM values across the full CBF/ATT range within the sample (g, h)

**Supplementary Table 1a:** Model 1 - Whole-WM MD/FA Relationships with Perfusion (Covaried for Sex). All models are multivariate linear regressions based on baseline-normalized measures (x_perc_) with overall F-statistics, *p*-values, and effect sizes (*R^2^_adjusted_*) listed. Slopes are noted below each regression equation and significant associations bolded with asterisks indicating significance. (*:p<.05, **:p<.01, ***:p<.001).

|  | **MD<sub>perc</sub>** |  | **FA<sub>perc</sub>** |  |
| --- | --- | --- | --- | --- |
| Tract | CBF<sub>perc</sub> | ATT<sub>perc</sub> | CBF<sub>perc</sub> | ATT<sub>perc</sub> |
| All | $F(2, 497) = 5.36, p = .005, R^2_{\text{Adjusted}} = .02$ | $F(2, 497) = 4.07, p = .02, R^2_{\text{Adjusted}} = .01$ | $F(2, 497) = 0.40, p = .67, R^2_{\text{Adjusted}} = -.002$ | $F(2, 497) = 0.02, p = .98, R^2_{\text{Adjusted}} = -.004$ |
|  | **-0.018508\*\*** | **0.231872\*** | 0.0089404 | 0.0060844 |

**Supplementary Table 1b:** Model 1 - Tractwise MD/FA Relationships with Perfusion (Covaried for Sex). All models are multivariate linear regressions based on baseline-normalized measures (x_perc_) with overall F-statistics, *p*-values, and effect sizes (*R^2^_adjusted_*) listed. Slopes are noted below each regression equation and significant associations bolded with asterisks indicating significance. (*:p<.05, **:p<.01, ***:p<.001).

|  | **MD<sub>perc</sub>** |  | **FA<sub>perc</sub>** |  |
| --- | --- | --- | --- | --- |
| Tract | CBF<sub>perc</sub> | ATT<sub>perc</sub> | CBF<sub>perc</sub> | ATT<sub>perc</sub> |
| Fmajor | $F(2, 490) = 7.47, p < .001, R^2_{\text{Adjusted}} = .03$ | $F(2, 490) = 5.68, p = .004, R^2_{\text{Adjusted}} = .02$ | $F(2, 490) = 8.24, p < .001, R^2_{\text{Adjusted}} = .03$ | $F(2, 490) = 8.08, p < .001, R^2_{\text{Adjusted}} = .03$ |
|  | -0.009599 | 0.081493 | -0.003605 | -0.010087 |
| Fminor | $F(2, 490) = 1.52, p = .22, R^2_{\text{Adjusted}} = .002$ | $F(2, 490) = 6.48, p = .002, R^2_{\text{Adjusted}} = .02$ | $F(2, 490) = 5.49, p = .004, R^2_{\text{Adjusted}} = .02$ | $F(2, 490) = 5.53, p = .004, R^2_{\text{Adjusted}} = .02$ |
|  | -0.011735 | **0.289540\*\*** | 0.019804 | -0.227797 |
| ATR | $F(2, 494) = 2.08, p = .13, R^2_{\text{Adjusted}} = .004$ | $F(2, 494) = 9.13, p < .001, R^2_{\text{Adjusted}} = .03$ | $F(2, 494) = 3.23, p = .009, R^2_{\text{Adjusted}} = .04$ | $F(2, 494) = 4.22, p = .015, R^2_{\text{Adjusted}} = .01$ |
|  | -0.014611 | **0.324941\*\*\*** | 0.027963 | **-0.306558\*\*** |
| CAB | $F(2, 487) = 0.47, p = .62, R^2_{\text{Adjusted}} = -.002$ | $F(2, 487) = 2.69, p = .07, R^2_{\text{Adjusted}} = .007$ | $F(2, 487) = 0.75, p = .47, R^2_{\text{Adjusted}} = -.001$ | $F(2, 487) = 4.53, p = .01, R^2_{\text{Adjusted}} = .01$ |
|  | 0.0006211 | -0.202510 | 0.01026 | **0.79903\*** |
| CCG | $F(2, 492) = 2.48, p = .085, R^2_{\text{Adjusted}} = .006$ | $F(2, 492) = 4.82, p = .008, R^2_{\text{Adjusted}} = .02$ | $F(2, 492) = 13.3, p < .001, R^2_{\text{Adjusted}} = .05$ | $F(2, 492) = 2.60, p = .006, R^2_{\text{Adjusted}} = .08$ |
|  | -0.013649 | **0.216401\*\*** | **0.068524\*\*\*** | 0.050215 |
| CST | $F(2, 488) = 6.42, p = .002, R^2_{\text{Adjusted}} = .02$ | $F(2, 488) = 6.19, p = .002, R^2_{\text{Adjusted}} = .02$ | $F(2, 488) = 1.89, p = .15, R^2_{\text{Adjusted}} = .004$ | $F(2, 488) = 1.80, p = .17, R^2_{\text{Adjusted}} = .003$ |
|  | -0.006782 | 0.050971 | -0.018407 | 0.225689 |
| ILF | $F(2, 494) = 8.03, p < .001, R^2_{\text{Adjusted}} = .03$ | $F(2, 494) = 3.67, p = .026, R^2_{\text{Adjusted}} = .01$ | $F(2, 494) = 2.28, p = .10, R^2_{\text{Adjusted}} = .005$ | $F(2, 494) = 2.90, p = .06, R^2_{\text{Adjusted}} = .008$ |
|  | **-0.019481\*** | -0.032828 | 0.004481 | 0.16351 |
| SLFP | $F(2, 494) = 3.64, p = .027, R^2_{\text{Adjusted}} = .01$ | $F(2, 494) = 5.51, p = .004, R^2_{\text{Adjusted}} = .02$ | $F(2, 494) = 2.15, p = .12, R^2_{\text{Adjusted}} = .005$ | $F(2, 494) = 4.46, p = .012, R^2_{\text{Adjusted}} = .01$ |
|  | -0.014721 | **0.214620\*** | -0.007393 | -0.247955 |
| SLFT | $F(2, 492) = 5.01, p = .007, R^2_{\text{Adjusted}} = .02$ | $F(2, 492) = 7.78, p = .008, R^2_{\text{Adjusted}} = .02$ | $F(2, 492) = 4.99, p = .007, R^2_{\text{Adjusted}} = .02$ | $F(2, 492) = 3.37, p = .035, R^2_{\text{Adjusted}} = .009$ |
|  | **-0.021370\*** | **0.222030\*** | -0.021391 | -0.127256 |
| UNC | $F(2, 492) = 0.49, p = .61, R^2_{\text{Adjusted}} = -.002$ | $F(2, 492) = 0.82, p = .44, R^2_{\text{Adjusted}} = -.0007$ | $F(2, 492) = 5.06, p = .007, R^2_{\text{Adjusted}} = .02$ | $F(2, 492) = 4.33, p = .014, R^2_{\text{Adjusted}} = .01$ |
|  | -0.0055602 | 0.1058895 | 0.015398 | -0.143507 |

**Supplementary Table 2a:** Model 2 - Whole-WM Sex Differences in Relationships Between MD/FA and Perfusion (Covaried for Age). All models are multivariate linear regressions based on baseline-normalized measures (x_perc_) with overall F-statistics, *p*-values, and effect sizes (*R^2^_adjusted_*) listed. Slopes are noted below each regression equation and significant associations bolded with asterisks indicating significance. (*:p<.05, **:p<.01, ***:p<.001).

|  | MD<sub>perc</sub> |  |  |  | FA<sub>perc</sub> |  |  |  |
| --- | --- | --- | --- | --- | --- | --- | --- | --- |
|  | CBF<sub>perc</sub> |  | ATT<sub>perc</sub> |  | CBF |  | ATT |  |
| Tract | Female | Male | Female | Male | Female | Male | Female | Male |
| All | $F(2, 291) = 55.5$,<br>$p < .001$, $R^2_{\text{Adjusted}} = .27$ | $F(2, 203) = 33.2$,<br>$p < .001$,<br>$R^2_{\text{Adjusted}} = .17$ | $F(2, 291) = 55.1$, $p < .001$,<br>$R^2_{\text{Adjusted}} = .27$ | $F(2, 203) = 22.3$,<br>$p < .001$,<br>$R^2_{\text{Adjusted}} = .17$ | $F(2, 291) = 9.72$, $p < .001$,<br>$R^2_{\text{Adjusted}} = .06$ | $F(2, 203) = 1.86$, $p = .16$,<br>$R^2_{\text{Adjusted}} = .008$ | $F(2, 291) = 9.51$,<br>$p < .001$, $R^2_{\text{Adjusted}} = .05$ | $F(2, 203) = 1.60$,<br>$p = .20$, $R^2_{\text{Adjusted}} = .006$ |
|  | -0.0079527 | -0.0007841 | 0.0559703 | -0.0668848 | 0.0130418 | -0.0143295 | 0.1261818 | 0.1857851 |

**Supplementary Table 2b:** Model 2 - Tractwise Sex Differences in Relationships Between MD/FA and Perfusion (Covaried for Age). All models are multivariate linear regressions based on baseline-normalized measures (x_perc_) with overall F-statistics, *p*-values, and effect sizes (*R^2^_adjusted_*) listed. Slopes are noted below each regression equation and significant associations bolded with asterisks indicating significance. (*:p<.05, **:p<.01, ***:p<.001).

|  | MD<sub>perc</sub> |  |  |  | FA<sub>perc</sub> |  |  |  |
| --- | --- | --- | --- | --- | --- | --- | --- | --- |
|  | CBF<sub>perc</sub> |  | ATT<sub>perc</sub> |  | CBF<sub>perc</sub> |  | ATT<sub>percentage</sub> |  |
| Tract | Female | Male | Female | Male | Female | Male | Female | Male |
| Fmajor | $F(2, 287) = 31.3$,<br>$p < .001$, $R^2_{\text{Adjusted}} = .17$ | $F(2, 200) = 9.30$,<br>$p < .001$,<br>$R^2_{\text{Adjusted}} = .009$ | $F(2, 287) = 32.9$, $p < .001$,<br>$R^2_{\text{Adjusted}} = .18$ | $F(2, 200) = 8.95$,<br>$p < .001$,<br>$R^2_{\text{Adjusted}} = .07$ | $F(2, 287) = 3.44$, $p = .034$,<br>$R^2_{\text{Adjusted}} = .02$ | $F(2, 200) = 1.96$, $p = .14$,<br>$R^2_{\text{Adjusted}} = .009$ | $F(2, 287) = 2.92$,<br>$p = .056$, $R^2_{\text{Adjusted}} = .01$ | $F(2, 200) = 1.96$,<br>$p = .14$, $R^2_{\text{Adjusted}} = .009$ |
|  | -0.0039667 | -0.0076777 | 0.1285074 | 0.0116263 | -0.0093200 | -0.0026277 | -0.0444465 | 0.0518249 |
| Fminor | $F(2, 288) = 77.5$,<br>$p < .001$, $R^2_{\text{Adjusted}} = .35$ | $F(2, 199) = 26.6$,<br>$p < .001$,<br>$R^2_{\text{Adjusted}} = .20$ | $F(2, 288) = 76.8$, $p < .001$,<br>$R^2_{\text{Adjusted}} = .34$ | $F(2, 199) = 26.7$,<br>$p < .001$,<br>$R^2_{\text{Adjusted}} = .20$ | $F(2, 288) = 24.4$, $p < .001$,<br>$R^2_{\text{Adjusted}} = .14$ | $F(2, 199) = 10.6$, $p < .001$,<br>$R^2_{\text{Adjusted}} = .09$ | $F(2, 288) = 22.0$,<br>$p < .001$, $R^2_{\text{Adjusted}} = .13$ | $F(2, 199) = 10.6$,<br>$p < .001$, $R^2_{\text{Adjusted}} = .09$ |
|  | -0.0093523 | -0.0011248 | 0.0490044 | 0.0443195 | 0.0248039 | 0.0045800 | -0.0178029 | -0.061765 |
| ATR | $F(2, 289) = 63.6$,<br>$p < .001$, $R^2_{\text{Adjusted}} = .30$ | $F(2, 202) = 32.7$,<br>$p < .001$,<br>$R^2_{\text{Adjusted}} = .24$ | $F(2, 289) = 65.0$, $p < .001$,<br>$R^2_{\text{Adjusted}} = .31$ | $F(2, 202) = 32.3$,<br>$p < .001$,<br>$R^2_{\text{Adjusted}} = .23$ | $F(2, 289) = 8.25$, $p < .001$,<br>$R^2_{\text{Adjusted}} = .05$ | $F(2, 202) = 0.25$, $p = .78$,<br>$R^2_{\text{Adjusted}} = -.007$ | $F(2, 289) = 8.44$,<br>$p < .001$, $R^2_{\text{Adjusted}} = .05$ | $F(2, 202) = 0.08$,<br>$p = .92$, $R^2_{\text{Adjusted}} = -.009$ |
|  | -0.0023397 | -0.0075814 | 0.1296512 | -0.0436909 | **0.0427019\*** | 0.0094317 | **-0.3579011\*** | -0.0112258 |
| CAB | $F(2, 285) = 14.5$,<br>$p < .001$, $R^2_{\text{Adjusted}} = .09$ | $F(2, 199) = 2.56$,<br>$p = .08$, $R^2_{\text{Adjusted}} = .02$ | $F(2, 285) = 14.3$, $p < .001$,<br>$R^2_{\text{Adjusted}} = .08$ | $F(2, 199) = 4.17$,<br>$p = .016$,<br>$R^2_{\text{Adjusted}} = .03$ | $F(2, 285) = 7.74$, $p < .001$,<br>$R^2_{\text{Adjusted}} = .04$ | $F(2, 199) = 4.04$, $p = .02$,<br>$R^2_{\text{Adjusted}} = .03$ | $F(2, 285) = 11.2$,<br>$p < .001$, $R^2_{\text{Adjusted}} = .07$ | $F(2, 199) = 4.68$,<br>$p = .01$, $R^2_{\text{Adjusted}} = .04$ |
|  | 0.0114287 | 0.0064030 | -0.1455464 | -0.3302063 | -0.0086050 | -0.0161303 | **0.7990257\*** | 0.7374741 |
| CCG | $F(2, 289) = 41.1$,<br>$p < .001$, $R^2_{\text{Adjusted}} = .22$ | $F(2, 200) = 18.6$,<br>$p < .001$,<br>$R^2_{\text{Adjusted}} = .15$ | $F(2, 289) = 42.5$, $p < .001$,<br>$R^2_{\text{Adjusted}} = .22$ | $F(2, 200) = 18.4$,<br>$p < .001$,<br>$R^2_{\text{Adjusted}} = .15$ | $F(2, 289) = 17.9$, $p < .001$,<br>$R^2_{\text{Adjusted}} = .10$ | $F(2, 200) = 11.5$, $p < .001$,<br>$R^2_{\text{Adjusted}} = .09$ | $F(2, 289) = 14.8$,<br>$p < .001$, $R^2_{\text{Adjusted}} = .09$ | $F(2, 200) = 6.26$,<br>$p = .002$, $R^2_{\text{Adjusted}} = .05$ |
|  | -0.010615 | -0.0046832 | 0.1605160 | -0.0354249 | **0.0482986\*** | **0.0755159\*\*\*** | 0.107982 | 0.5138831 |
| CST | $F(2, 284) = 9.29$,<br>$p < .001$, $R^2_{\text{Adjusted}} = .05$ | $F(2, 201) = 1.37$,<br>$p = .26$, $R^2_{\text{Adjusted}} = .004$ | $F(2, 284) = 9.60$, $p < .001$,<br>$R^2_{\text{Adjusted}} = .06$ | $F(2, 201) = 1.35$,<br>$p = .26$, $R^2_{\text{Adjusted}} = .003$ | $F(2, 284) = 0.97$, $p = .38$,<br>$R^2_{\text{Adjusted}} = -.0002$ | $F(2, 201) = 3.67$, $p = .03$,<br>$R^2_{\text{Adjusted}} = .03$ | $F(2, 284) = 1.97$,<br>$p = .14$, $R^2_{\text{Adjusted}} = .007$ | $F(2, 201) = 1.29$,<br>$p = .28$, $R^2_{\text{Adjusted}} = .003$ |
|  | 0.0061903 | -0.0065296 | -0.1043070 | 0.0842141 | -0.0054017 | **-0.03705\*\*** | 0.2295747 | 0.3193 |
| ILF | $F(2, 289) = 43.0$,<br>$p < .001$, $R^2_{\text{Adjusted}} = .22$ | $F(2, 202) = 17.7$,<br>$p < .001$,<br>$R^2_{\text{Adjusted}} = .14$ | $F(2, 289) = 41.8$, $p < .001$,<br>$R^2_{\text{Adjusted}} = .22$ | $F(2, 202) = 17.7$,<br>$p < .001$,<br>$R^2_{\text{Adjusted}} = .14$ | $F(2, 289) = 10.3$, $p < .001$,<br>$R^2_{\text{Adjusted}} = .06$ | $F(2, 202) = 3.82$, $p = .02$,<br>$R^2_{\text{Adjusted}} = .03$ | $F(2, 289) = 10.7$,<br>$p < .001$, $R^2_{\text{Adjusted}} = .06$ | $F(2, 202) = 3.50$,<br>$p = .03$, $R^2_{\text{Adjusted}} = .02$ |
|  | -0.0125893 | 0.0018092 | 0.0240348 | 0.0051664 | -0.0020199 | -0.0112217 | 0.1423583 | 0.0884416 |
| SLFP | $F(2, 288) = 47.8$,<br>$p < .001$, $R^2_{\text{Adjusted}} = .24$ | $F(2, 203) = 20.6$,<br>$p < .001$,<br>$R^2_{\text{Adjusted}} = .16$ | $F(2, 288) = 48.1$, $p < .001$,<br>$R^2_{\text{Adjusted}} = .25$ | $F(2, 203) = 20.8$,<br>$p < .001$,<br>$R^2_{\text{Adjusted}} = .16$ | $F(2, 288) = 7.54$, $p < .001$,<br>$R^2_{\text{Adjusted}} = .04$ | $F(2, 203) = 1.73$, $p = .18$,<br>$R^2_{\text{Adjusted}} = .007$ | $F(2, 288) = 8.76$,<br>$p < .001$, $R^2_{\text{Adjusted}} = .05$ | $F(2, 203) = 0.41$,<br>$p = .67$, $R^2_{\text{Adjusted}} = -.006$ |
|  | -0.0059281 | 0.0023829 | 0.0836243 | 0.0632643 | -0.00005828 | -0.02840 | -0.2094319 | -0.1595419 |
| SLFT | $F(2, 287) = 52.9$,<br>$p < .001$, $R^2_{\text{Adjusted}} = .26$ | $F(2, 202) = 22.2$,<br>$p < .001$,<br>$R^2_{\text{Adjusted}} = .17$ | $F(2, 287) = 53.6$, $p < .001$,<br>$R^2_{\text{Adjusted}} = .27$ | $F(2, 202) = 22.3$,<br>$p < .001$,<br>$R^2_{\text{Adjusted}} = .17$ | $F(2, 287) = 4.30$, $p = .014$,<br>$R^2_{\text{Adjusted}} = .02$ | $F(2, 202) = 6.02$, $p = .003$,<br>$R^2_{\text{Adjusted}} = .05$ | $F(2, 287) = 4.59$,<br>$p = .011$, $R^2_{\text{Adjusted}} = .02$ | $F(2, 202) = 1.80$,<br>$p = .17$, $R^2_{\text{Adjusted}} = .008$ |
|  | -0.0078787 | 0.0006048 | 0.1188965 | 0.0478854 | -0.0097048 | **-0.0440165\*\*** | -0.1308776 | -0.0510914 |
| UNC | $F(2, 290) = 49.5$,<br>$p < .001$, $R^2_{\text{Adjusted}} = .25$ | $F(2, 199) = 18.4$,<br>$p < .001$,<br>$R^2_{\text{Adjusted}} = .15$ | $F(2, 290) = 49.8$, $p < .001$,<br>$R^2_{\text{Adjusted}} = .25$ | $F(2, 199) = 18.4$,<br>$p < .001$,<br>$R^2_{\text{Adjusted}} = .15$ | $F(2, 290) = 3.21$, $p = .04$,<br>$R^2_{\text{Adjusted}} = .01$ | $F(2, 199) = 0.30$, $p = .74$,<br>$R^2_{\text{Adjusted}} = -.007$ | $F(2, 290) = 1.28$,<br>$p = .28$, $R^2_{\text{Adjusted}} = .002$ | $F(2, 199) = 0.32$,<br>$p = .73$, $R^2_{\text{Adjusted}} = -.007$ |
|  | 0.00004331 | 0.0009167 | 0.0632672 | -0.0315701 | 0.0289662 | -0.0053114 | -0.1422849 | -0.0935715 |

